# Neurochondrin interacts with the SMN protein suggesting a novel mechanism for Spinal Muscular Atrophy pathology

**DOI:** 10.1101/183640

**Authors:** Luke Thompson, Kim Morrison, Sally Shirran, Catherine Botting, Judith Sleeman

## Abstract

Spinal Muscular Atrophy (SMA) is an inherited neurodegenerative condition caused by reduction in functional Survival Motor Neurones Protein (SMN). SMN has been implicated in transport of mRNA in neural cells for local translation. We previously identified microtubule-dependant mobile vesicles rich in SMN and the splicing factor SmB, a member of the Sm protein family, in neural cells. By comparing the proteome of SmB to that of SmN, a neural-specific Sm protein, we now show that the essential neural protein neurochondrin (NCDN) interacts with Sm proteins and SMN in the context of mobile vesicles in neurites. NCDN has roles in protein localisation in neural cells, and in maintenance of cell polarity. NCDN is required for the correct localisation of SMN, suggesting they may both be required for formation and transport of trafficking vesicles. NCDN provides a potential therapeutic target for SMA together with, or in place of, those targeting SMN expression.

## Introduction

The inherited neurodegenerative disease, Spinal Muscular Atrophy (SMA) is caused by a reduction in the amount of functional Survival Motor Neuron (SMN) protein (Lefebvre et al. 1995). SMA is the leading genetic cause of infant mortality, affecting 1:6000 live births (Monani 2005). The recently developed therapy, Spinraza/Nusinersin (Biogen) has been shown to increase the level of SMN and improve the symptoms of SMA patients (Passini et al. 2011, Finkel et al. 2016, Corey 2017). Most SMA patients harbour mutations in the SMN1 gene, which produces the majority of total SMN protein in cells. In humans, expression from a variable number of copies of an additional gene, SMN2, can produce some full-length SMN protein (Lefebvre et al. 1995, Lefebvre et al. 1997). The SMN2 gene, unlike SMN1, contains a point mutation in an exon splicing enhancer (Lorson et al. 1999, Lorson, Androphy 2000) resulting in truncation of most of the SMN protein produced by SMN2 through skipping of exon 7. The truncated protein produced by SMN2 is less stable than full-length SMN and cannot compensate fully for the loss of SMN1 (Lorson et al. 1999, Lorson, Androphy 2000). However, due to the small amounts of full length SMN expressed from the SMN2 gene, the number of gene copies can influence the severity of SMA, with evidence that five copies of SMN2 may be enough to compensate for loss of SMN1 (Campbell et al. 1997, Prior et al. 2004). It is not currently clear how a deficiency of functional SMN leads to the specific symptoms of SMA. In particular, the differing sensitivity of cell types to lowered SMN levels, with motor neurons most severely affected, is difficult to explain as SMN is an essential protein and complete deletion is lethal at the cellular level (Schrank et al. 1997, Hsieh-Li et al. 2000).

SMN localises to nuclear bodies called gems (Gemini of Cajal bodies (CBs)) (Liu, Dreyfuss 1996) as well as in the cytoplasm where it has two well-established cellular roles (Monani 2005, Sleeman 2013, Li et al. 2014, Tisdale, Pellizzoni 2015). The first to be elucidated was a role in the early stages of assembly and maturation of splicing snRNPs (small nuclear ribonucleoproteins). Splicing snRNPs are ribonucleoprotein complexes, essential for pre-mRNA splicing, comprising an snRNA (small nuclear RNA) core and numerous proteins including a heptameric ring containing one copy each of members of the Sm protein family. SMN is part of a cytoplasmic complex, also containing the gemin proteins, required for the addition the Sm proteins as a ring around the snRNA core (Liu et al. 1997, Stark et al. 2001, Li et al. 2014, Tisdale, Pellizzoni 2015). The maturation of snRNPs has been shown to be impaired by a deletion in SMN (Shpargel, Matera 2005, Wan et al. 2005, Winkler et al. 2005, Gabanella et al. 2007, Zhang et al. 2008), while alterations to pre-mRNA splicing events, proposed to be a downstream consequence of this impairment, have been observed in several models of SMA (Zhang et al. 2008, Custer et al. 2013, Huo et al. 2014). One of the proposed mechanisms for the cell-type specificity of SMA is that these alterations of pre-mRNA splicing events affect mRNA transcripts that are essential for motor neurons, perhaps preferentially affecting transcripts spliced by the minor spliceosome (Gabanella et al. 2007, Zhang et al. 2008, Boulisfane et al. 2011, Custer et al. 2016, Doktor et al. 2017). Despite promising results in Drosophila models, however, specific transcripts affecting MNs are yet to be conclusively identified (Lotti et al. 2012).

The second established cellular role of SMN is in the trafficking of mature mRNA within the cytoplasm, particularly in the axons and neurites of neural cell types. (Rossoll et al. 2002, Rossoll et al. 2003a, Zhang et al. 2003, Zhang et al. 2006a, Todd et al. 2010a, Todd et al. 2010b, Akten et al. 2011, Fallini et al. 2011, Peter et al. 2011, Custer et al. 2013, Fallini et al. 2014, Li et al. 2015, Fallini et al. 2016). This is thought to be linked to local translation of mRNA into proteins, an important process for neural cells, in particular motor neurones, due to the length of their axons (Kang, Schuman 1996, Huber et al. 2000, Doyle, Kiebler 2011, Holt, Schuman 2013), making this trafficking role for SMN of particular interest for understanding the cellular pathology of SMA. SMN co-localises with the mRNA binding proteins HuD, IMP1 and hnRNP R and is involved in the localisation of mRNAs to axons (Rossoll et al. 2003b, Fallini et al. 2011, Akten et al. 2011, Fallini et al. 2014). The cellular structures involved in SMN-dependent mRNA trafficking are currently unclear, being described as granular (Zhang et al. 2003, Peter et al. 2011, Fallini et al. 2014, Akten et al. 2011, Zhang et al. 2006b, Todd et al. 2010b) or vesicular in nature (Custer et al. 2013, Prescott et al. 2014).

Structures containing the SMN and SmB proteins, alongside Coatomer Gamma are trafficked on microtubules (Prescott et al. 2014). Evidence suggesting that SMN/SmB-rich structures are vesicular in nature includes their staining with lipophilic dyes in living neural cells, their vesicular appearance using correlative fluorescence and electron microscopy (Prescott et al. 2014), and the interaction of SMN and the Sm proteins with Coatomer proteins, which are associated with membrane bound vesicles (Peter et al. 2011, Custer et al. 2013, Prescott et al. 2014). We have previously identified an interaction between SmB and dynein cytoplasmic 1 heavy chain 1 (DYNC1H1), a motor protein required for microtubule transport (Prescott et al. 2014), mutation in which can cause a rare, lower extremity dominant SMA (Harms et al. 2010, Harms et al. 2012, Tsurusaki et al. 2012, Niu et al. 2015, Peeters et al. 2015, Punetha et al. 2015, Scoto et al. 2015, Strickland et al. 2015, Ding et al. 2016, Chen et al. 2017). SMN has also been shown to associate with the membranous Golgi complex (Ting et al. 2012), while mutations in the Golgi-related protein BICD2 (bicaudal D homolog 2) causing a form of lower extremity dominant SMA (Neveling et al. 2013, Oates et al. 2013, Peeters et al. 2013, Synofzik et al. 2014, Martinez-Carrera, Wirth 2015, Rossor et al. 2015).

The Sm protein family is implicated in both of the major functions of SMN. The core of splicing snRNPs comprises a heptameric ring of proteins around the snRNA, containing SmB/B', D1, D2, D3 E, F and G (Urlaub et al. 2001). SMN is a vital part of the complex required for the assembly of this Sm protein ring (Liu, Dreyfuss 1996, Fischer et al. 1997, Meister et al. 2001, Meister, Fischer 2002, Pellizzoni et al. 2002, Battle et al. 2006, Fischer et al. 2011). Other members of the Sm protein family have also been identified, beyond the core proteins usually found in splicing snRNPs. Of particular interest in the context of SMA pathology is SmN (encoded by the SNRPN gene), which is expressed in neural tissues (Schmauss et al. 1992) and can replace SmB in the heptameric Sm protein ring (Huntriss et al. 1993). The human SmN protein differs from SmB' by 17 amino acids (UniProt Identifiers P63162 and P14678 respectively), and little is known about its behaviour other than its incorporation into snRNPs, although the SNRPN gene locus is within the paternally imprinted region of the genome critical in Prader-Willi Syndrome (Ozcelik et al. 1992). There is growing appreciation that some Sm proteins may ‘moonlight' in functions beyond their presence in splicing snRNPs. In addition to the role of SmB in cytoplasmic trafficking vesicles in human cells, in Drosophila SmB and SmD3 are implicated in mRNA localisation (Gonsalvez et al. 2010) and SmD1 has a role in miRNA biogenesis (Xiong et al. 2015). With this in mind, we applied a proteomic approach in the neural cell line SH-SY5Y to search for interactions that could indicate neural specific roles for the SmN protein of relevance for the pathology of SMA.

Our proteomic approach led to the identification of a novel interaction between SmN and the essential protein Neurochondrin (NCDN). NCDN is predominantly expressed in neural tissue, where it localises to dendrites, and is involved in the promotion of neural outgrowth (Shinozaki et al. 1997, Shinozaki et al. 1999, Wang et al. 2013, Oku et al. 2013) and the regulation of synaptic plasticity (Dateki et al. 2005, Wang et al. 2009). At a molecular level, NCDN is implicated in the moderation of several signalling pathways in neural cells (Dateki et al. 2005, Francke et al. 2006, Wang et al. 2009, Ward et al. 2009, Wang et al. 2013, Matosin et al. 2015, Pan et al. 2016). NCDN may also have a function in cell polarity maintenance, within dendritic Rab5 positive vesicles (Oku et al. 2013, Guo et al. 2016). Further investigation of NCDN in the context of SMA pathology revealed that it also interacts with SMN. Furthermore, NCDN appears to be required for the correct sub-cellular localisation of SMN. These data suggest that NCDN is involved in a novel, neural-specific role for SMN in the context of mobile cytoplasmic vesicles that also contain a subset of the Sm protein family. This has implications both for understanding the cellular pathology of SMA and for the development of additional therapies for the condition.

## Results

### SmN exhibits similar behaviour to SmB, localising to cytoplasmic vesicles containing SMN

In order to determine the interactome of the neural-specific Sm protein, SmN, we first generated constructs to express fluorescent protein-tagged SmN by amplifying the SmN sequence from total RNA from SH-SY5Y cells and cloning it into the pEYFP-C1 and pmCherry-C1vectors. All of the Sm proteins studied so far show a steady-state localisation to nuclear Cajal Bodies (CBs) and speckles. The Sm proteins SmB, SmD1 and SmE have also previously been shown to exhibit a characteristic pathway within the cell on their initial expression, indicative of the snRNP maturation pathway (Sleeman, Lamond 1999), although differences were seen between the Sm proteins. To confirm that YFP-SmN localised correctly at steady state to CBs and speckles, and to determine where SmN localised during maturation and incorporation into snRNPs, the plasmid was transiently expressed in SH-SY5Y neuroblastoma cells, with cells fixed and immunostained at 24 hour intervals. At 48 and 72 hours after transfection, YFP-SmN predominantly localised to speckles and Cajal bodies (CBs, detected with anti-coilin) identically to endogenous Sm proteins (detected with Y12 antibody), whereas at 24 hours, YFP-SmN localised predominantly diffusely within the cytoplasm, with some accumulation in CBs (Fig 1A, B). This sequential localisation is indistinguishable from that previously observed with YFP-SmB In HeLa and MCF-7 cells, though CBs were not prominent in the majority of SH-SY5Y cells transiently expressing YFP-SmN. Equivalent results were obtained in SH-SY5Y cells transiently expressing mCherry-SmN (Fig S1). Both YFP-SmN and mCherry-SmN are efficiently incorporated into splicing snRNPs, as evidenced by their enrichment from whole-cell lysates using antibodies against the characteristic hypermethylated Cap structure (2,2,7-trimethylaguanosine) found on snRNAs (Fig 1C).

**Figure 1:**
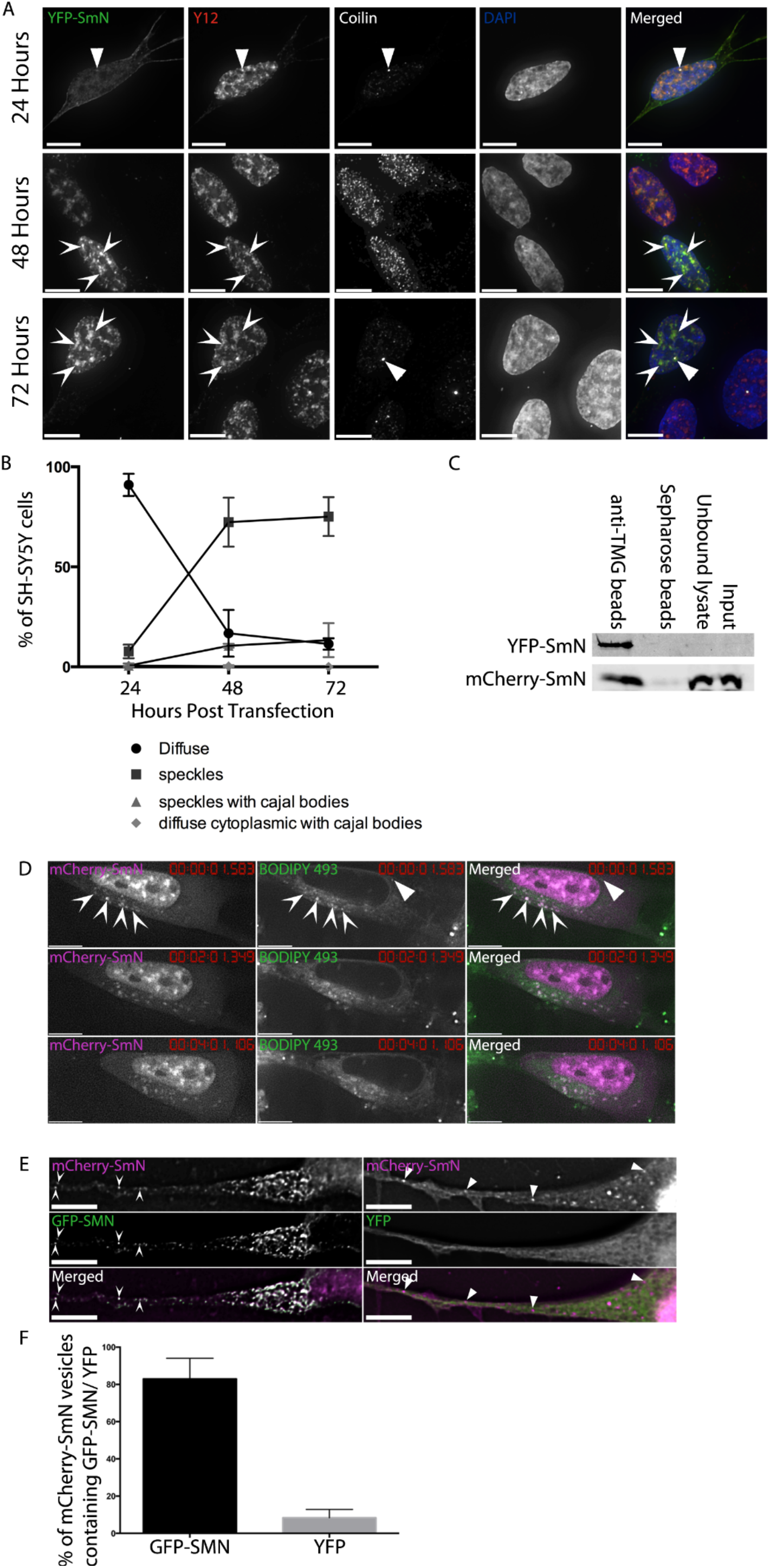
SmN exhibits similar behaviour to SmB in SH-SY5Y cells. *A) SH-SY5Y cells transiently expressing YFP-SmN and fixed after 24, 48 and 72 hours show variations in distribution of the YFP-SmN with time. Immunostaining with Y12 (red on overlay) and anti-coilin (white on overlay) shows splicing speckles (arrowheads) and Cajal Bodies (CBs, triangles) respectively. Images are deconvolved z-stacks with 0.2 μm spacing. Bar=7μm. B) SmN initially localises diffusely in the cytoplasm, before localising to speckles at the 48 and 72 hour time-points. 3 independent experiments, n=100 cells per experiment. Data shown is mean +/– SD). C) Western blot analysis of snRNPs immunprecipiated using TMG beads (left hand lane) confirms that both YFP-SmN (detected with anti-YFP, top row) and mCherry-SmN (detected with anti-mCherry, bottom row) are incorporated into snRNPs. D) mCherry-SmN cytoplasmic structures are mobile and stain with the lipophilic dye BODIPY 493. Arrows identify mCherry-SmN structures stained with BODIPY 493, triangles identify BODIPY 493-stained vesicles not containing mCherry-SmN. Cells were imaged approximately every 4 seconds for 9 minutes. Images are single deconvolved z-sections. Bar=7μm. E) mCherry-SmN and GFP-SMN co-localise in cytoplasmic foci in SH-SY5Y cells (arrowheads in left hand panels), whereas YFP alone shows no accumulation in mCherry-SmN foci (triangles in right hand panels). White signal on the overlay indicates areas of co-localisation. Images are single deconvolved z-sections. Bar=7μm. F) Comparison of the percentage of mCherry-SmN vesicles per cell co-localising with GFP-SMN to those showing co-incidental overlap with YFP alone confirms the co-localisation (unpaired 2 tailed t-test, p=<0.0001, n=5).*

To determine whether the similarities between SmN and SmB extend to localisation in vesicles in the cytoplasm (Prescott et al. 2014), SH-SY5Y cells constitutively expressing mCherry-SmN were used for live-cell time-lapse microscopy. Mobile mCherry-SmN foci were observed (movie S1). In common with the SmB vesicles, these stained positive with the lipophilic dye, BODIPY 493, indicating that they are vesicular in nature (Fig 1D). Finally, to confirm that the mCherry-SmN vesicles were similar to those previously identified with SmB, SH-SY5Y cells constitutively expressing mCherry-SmN were transfected with plasmids to express either GFP-SMN or YFP alone. GFP-SMN co-localised with mCherry-SmN in 83% (±11) of SmN-positive vesicles, which is statistically significant when compared to 8.4% (±4.5) of mCherry-SmN vesicles co localising with YFP alone (Fig 1E, F).

### Mass Spectrometry reveals the similarities between the interactomes of SmN and SmB

As SmN appeared to behave very similarly to SmB in neural cells, it was unclear why neural cells express two almost identical proteins. It was decided to investigate whether SmN and SmB may have differing roles that could be identified by proteomic analysis. Proteins interacting with YFP-SmB and YFP-SmN were affinity purified using GFP-TRAP (Chromotek) from whole-cell lysates of SH-SY5Y cell lines constitutively expressing the tagged proteins, with a cell line expressing YFP alone as a control for non-specific binding to the tag or bead matrix. Immunoblot analysis using antibodies to YFP demonstrated that the enrichment of the tagged proteins was 20X, 23X and 4X for YFP-SmN, YFP-SmB and YFP alone respectively (Fig 2A). The affinity purified material was size separated using SDS-PAGE and analysed by nLC ESI MS/MS mass spectrometry to identify peptides and therefore proteins interacting with YFP-SmB and YFP-SmN. Following removal of likely contaminants identified by their interaction with YFP alone, or their previous identification as common interactors of GFP-TRAP (Trinkle-Mulcahy et al. 2008), UniProt Biological Process and Cellular Component Genome Ontology annotations were used to group identified proteins into categories depending on function. These groups were then used to determine whether there were differences in possible functions between SmN and SmB (Fig 2B). Numerous proteins previously established to interact with Sm proteins were identified including SMN and the gemins as well as the methylosome components PRMT5, MEP50 and pICIn, validating our approach (Table S2). The overall proportions of proteins in each category were similar when comparing the interactomes of SmN to SmB, though differences were identified at the level of individual proteins. Of particular interest in the context of SMA were a number of proteins with potential neural specific roles, which were identified in one or both samples. One of these was Neurochondrin (NCDN), a relatively poorly characterised neural protein, which was identified in the YFP-SmN interactome, with 5 unique peptides identified (Fig 2C).

**Figure 2:**
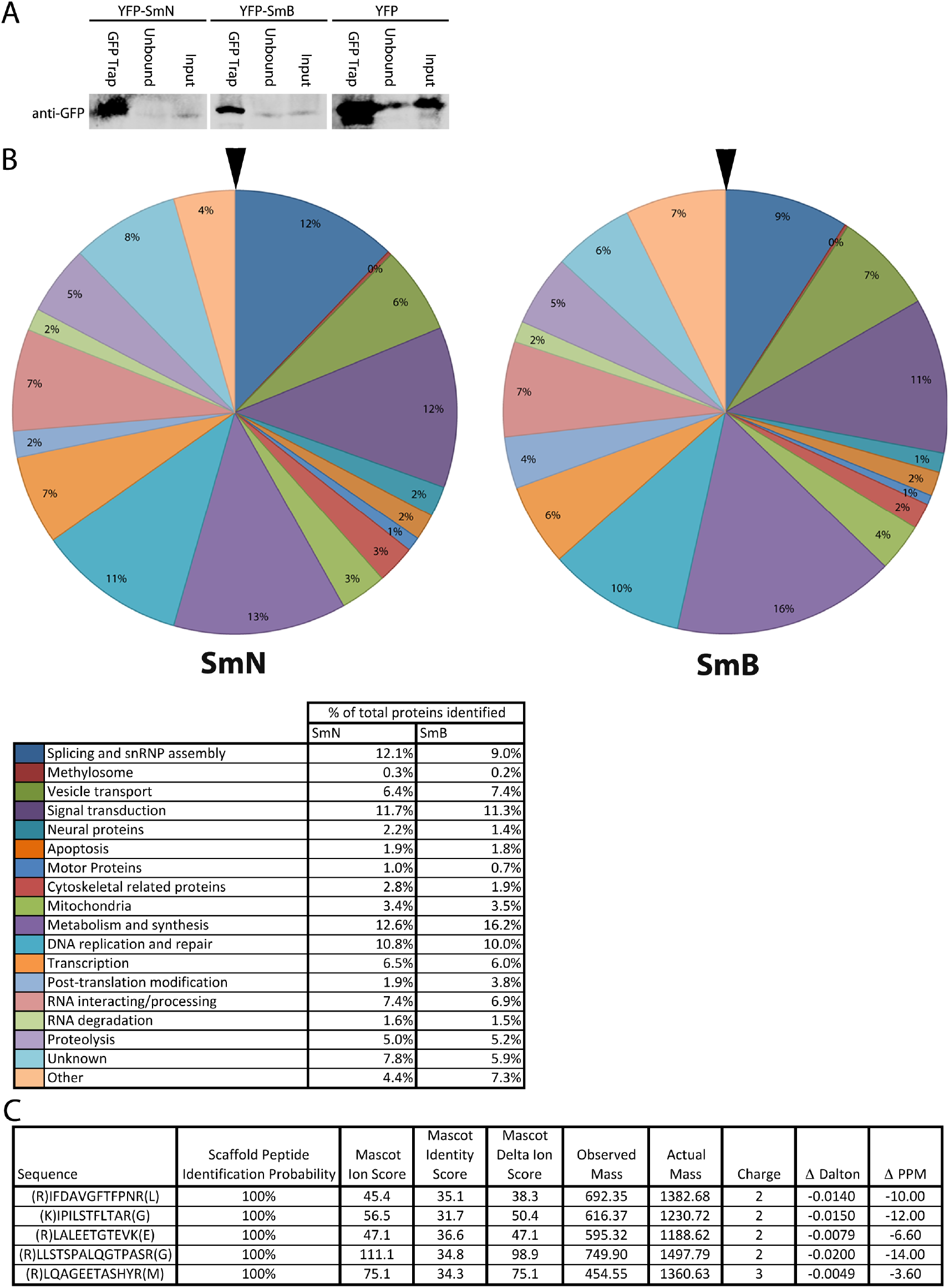
The interactomes of SmB and SmN are similar, but there are differences present at the level of individual proteins. *A) Immunoblot analysis confirms efficient affinity purification of YFP-SmN, YFP-SmB and YFP. 10% of the affinity purified material (left hand lane in each panel) was compared to 80μg of precleared lysate (Input) and unbound material using anti-GFP. GFP-Trap effectively immunoprecipitated all three proteins. B) After processing the mass spectrometry data, and sorting identified proteins into groups based on Gene Ontology annotations, the interactomes of SmN and SmB are very similar. C) Neurochondrin (NCDN) was identified in the interactome of YFP-SmN, with 5 unique peptide hits encompassing 9% sequence coverage. Each Ion score (Mascot Ion Score) was above the threshold for peptide identity (Mascot Identity Score), with 2 out of the 5 identified peptides having a score of above double the threshold score.*

### Neurochondrin interacts with SmN, SmB and SMN

To verify the interaction between Sm proteins and Neurochondrin, a construct expressing NCDN-GFP was generated. Affinity purification of NCDN-GFP from whole cell lysates of SH-SY5Y cells transiently co-expressing NCDN-GFP and mCherry-SmB demonstrated interaction between NCDN-GFP and mCherry-SmB (Fig 3A). To further investigate interactions between NCDN and the Sm proteins neural cells, an SH-SY5Y cell line constitutively expressing NCDN-GFP was established. Affinity purification of NCDN-GFP from whole cell lysates followed by immunoblot analysis using antibodies against endogenous SmN and SmB (Fig 3B) revealed that NCDN-GFP interacts with both SmN and SmB. Furthermore, both endogenous SMN and endogenous βCOP (a coatomer vesicle protein) were also revealed to interact with NCDN-GFP, suggesting that NCDN interacts with the Sm proteins and SMN in the context of cytoplasmic vesicles. Affinity purification of YFP alone from whole cell lysates of an SH-SY5Y cell line constitutively expressing YFP does not result in co purification of endogenous SMN, SmB or βCOP (Fig 3C). To further investigate the interaction between SMN and NCDN observed in Fig 3B, a reciprocal experiment was performed using GFP-Trap to affinity purify GFP-SMN from an SH-SY5Y cell line constitutively expressing GFP-SMN (Clelland et al. 2009). Subsequent immunoblot analysis using antibodies to endogenous NCDN demonstrated that NCDN co-enriched with GFP-SMN (Fig 3D). To determine the probable cellular location for the interaction between SmN/SmB and NCDN plasmids to express either NCDN-GFP or YFP were transiently transfected into SH-SY5Y cells constitutively expressing mCherry-SmN. This revealed that NCDN-GFP, but not YFP alone, accumulates in cytoplasmic vesicles containing mCherry-SmN (Fig3E). Similar results were obtained when NCDN-GFP was transiently expressed in SH-SY5Y cells constitutively expressing mCherry-SmB (Fig S3). To investigate the probable cellular location of interactions between SMN and NCDN, mCherry-SMN was co-expressed with either NCDN-GFP or YFP alone. NCDN-GFP was found in cytoplasmic structures enriched in mCherry-SMN (Fig 3F). These data suggest that NCDN co-localises with both the Sm proteins and SMN in cytoplasmic vesicles, although NCDN-GFP shows an increase diffuse signal compared to the Sm proteins or SMN.

**Figure 3:**
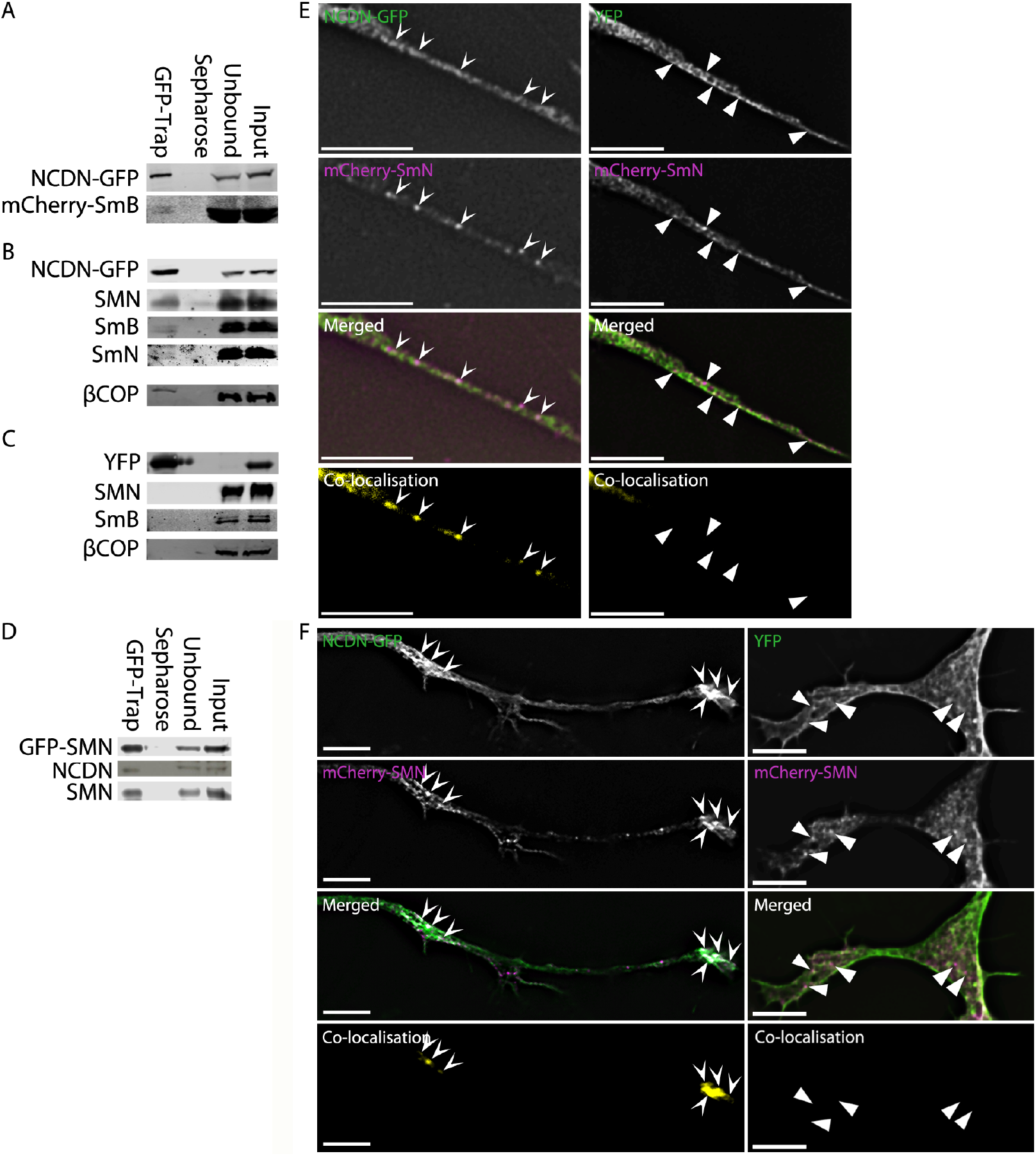
NCDN interacts with SmN, SmB and SMN, and co-localises with them in vesicles. *A) Affinity isolation of NCDN-GFP using GFP-Trap, detected with anti-GFP (top row) co-enriches mCherry-SmB, detected with anti-mCherry (bottom row) in transiently co-transfected SH-SY5Y cells B) In an SH-SY5Y cell line constitutively expressing NCDN-GFP, affinity isolation of NCDN-GFP, detected with anti-GFP (top row) co-enriches SMN, SmB, SmN and the Coatomer protein βCOP, all detected with antibodies against the endogenous proteins (as labelled). C) In an SH-SY5Y cell line constitutively expressing YFP, affinity isolation of YFP, detected with anti-GFP (top row) does not co-enrich SMN, SmB or βCOP, all detected with antibodies to the endogenous proteins (as labelled). D) in an SH-SY5Y cell line constitutively expressing GFP-SMN, affinity isolation of GFP-SMN, detected with anti-GFP (top row) co-enriches NCDN, detected with anti-NCDN (middle row). Endogenous SMN, detected with anti-SMN (bottom row) is also co-enriched. E) NCDN-GFP and mCherry-SmN co-localise in vesicle-like structures (arrows) in neurites of SH-SY5Y cells constitutively expressing mCherry-SmN, and transiently expressing NCDN-GFP (left hand panels). White areas in the merged image show areas of co-localisation. Co-localisation images (bottom row) were generated in Volocity, using automatic thresholds on undeconvolved z-sections before excluding values below 0.05 (see material and methods). No co-localisation is seen in the same cell line transiently expressing YFP alone (right hand panel). Triangles show structures containing mCherry-SmN, but not YFP. F) mCherry-SMN and NCDN-GFP co-localise in vesicles (arrows) in the cytoplasm of co-transfected SH-SY5Y cells (left hand panels). White areas in the merged image show areas of co-localisation. Co-localisation images (bottom row) were generated as above. No co-localisation is observed between mCherry-SMN and YFP (triangles, right hand panels). Bar= 7μm.*

### NCDN, SMN and Sm protein co-fractionate with coatomer proteins

To further investigate the possibility that the interaction of NCDN with SMN and the Sm proteins occurs within cytoplasmic vesicles, sub-cellular fractionations were performed on both parental SH-SY5Y cells and SH-SY5Y cell lines constitutively expressing NCDN-GFP, YFP-SmB, YFP-SmN or YFP. Sequential centrifugation was used to separate the cells into a nuclear fraction, 16,000 RCF and 100,000 RCF cytoplasmic pellets and cytosolic supernatant (de Duve 1971, Lodish et al. 2000, Mohr, Völkl 2001, Andersen et al. 2002). Immunoblotting of these sub-cellular fractions revealed that GFP-NCDN, YFP-SmB and YFP-SmN, were all enriched in the 100,000 RCF pellet, along with endogenous SMN and coatomer proteins (Fig 4). This fraction would be expected to contain membrane-bound structures, such as microsomes and small cytoplasmic vesicles, which would encompass small coatomer type endocytic vesicles (de Duve 1971, Mohr, Völkl 2001, Lodish et al. 2000). Endogenous SmN and SmB were also observed to enrich similarly (Fig S4). NCDN-GFP shows a larger proportion of protein in the remaining cytosolic supernatant when compared to YFP-SmB, YFP-SmN and endogenous SMN, which is in agreement with the subcellular localisations observed (Fig 3). This further supports our hypothesis that the interactions between NCDN, SMN and the Sm proteins take place in small cytoplasmic vesicles.

**Figure 4:**
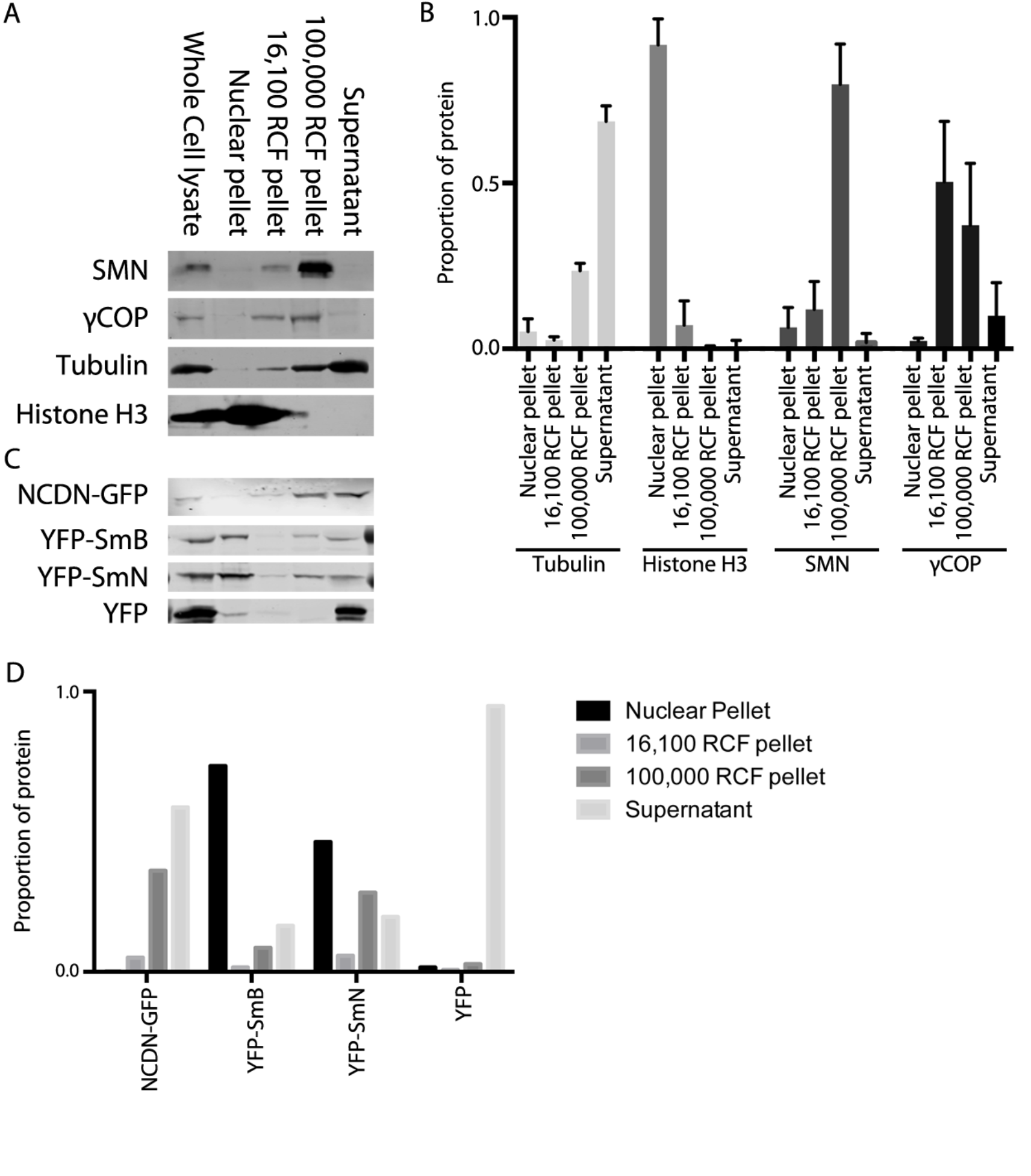
Detergent-free fractionation of SH-SY5Y cells reveals that SMN, coatomer proteins, NCDN, SmB and SmN are all enriched in the 100,000 RCF vesicle pellet. *A) Immunoblotting of equal protein amounts from fractionated SH-SY5Y cells reveals that SMN (top row) is highly enriched in the 100,000 RCF pellet, with smaller amounts seen in the 16,100 RCF pellet and the nuclear pellet. The coatomer protein, γCOP (second row) is also enriched in the 100,00 RCF pellet as well as the 16,100 RCF pellet. Antibodies to histone H3 and tubulin confirm minimal nuclear contamination in cytoplasmic fractions, and minimal cytoplasmic contamination in the nuclear pellet, respectively. B) Quantitation of immunoblot analysis confirms that SMN is highly enriched in the 100,000 RCF pellet, with enrichment of γCop also seen. Histone H3 and tubulin are highly enriched in the nucleus and cytoplasm respectively. Quantitation of tubulin and histone H3 bands was from 7 immunoblots, with values from SMN and γCOP from 5 and 4 immunoblots respectively. C) Immunoblotting of equal protein amounts from fractionated SH-SY5Y cells constitutively expressing NCDN-GFP, YFP-SmB, YFP-SmN or YFP alone (all detected with anti-GFP) reveals that NCDN-GFP is enriched in the 100,000 RCF pellet, with smaller amounts seen in the 16,100 RCF pellet and the cytosolic supernatant. YFP-SmB and YFP-SmN are both also found in the 100,000 RCF pellet, in addition to the nuclear pellet and cytosolic supernatant. YFP alone is found almost exclusively in the cytosolic supernatant, with none detected in the 100,000 RCF or 16,100 RCF pellets. D) Quantitation of the immunoblots in C) confirms the presence of NCDN-GFP, YFP-SmB and YFP-SmN in the 100,000 RCF pellet, together with the restriction of YFP alone to the residual cytosolic supernatant.*

### NCDN is required for the correct sub-cellular localisation of SMN

We have previously documented that reduction of SmB expression results in re-localisation of SMN into numerous nuclear structures, probably analogous to gems (Gemini of CBs), and its loss from cytoplasmic structures (Prescott et al. 2014). To investigate the requirement for NCDN in cytoplasmic SMN localisation, an SH-SY5Y cell line constitutively expressing GFP-SMN (Clelland et al. 2009) was transfected with siRNAs targeting NCDN (4 different single siRNAs (Dharmacon) and a pooled sample). A reduction in NCDN expression caused an increase in the number of SMN-positive gems present in the cell nucleus, as did a reduction of SmB expression (Fig 5A, B). Conversely, reduction in SMN expression reduced the number of nuclear gems. The use of non targeting control (siControl) sequences or positive control siRNAs (targeting Lamin A/C) had no effect on the number of nuclear gems. The reduction in gene expression, assayed by immunoblotting, for each siRNA was typically 40-60% (Fig 5C). This suggests that NCDN is required for the correct sub-localisation of SMN. Of potential relevance for SMA pathology, depletion of either NCDN or SmB causes GFP-SMN to adopt a sub-cellular localisation reminiscent of that shown by GFP-SMN∆7 (Fig 5), a truncated version of SMN that mimics the product of the SMN2 gene and is unable to substitute for full-length SMN in models of SMA (Monani et al. 1999, Monani et al. 2000).

### SMN is required for the correct sub-cellular localisation of NCDN

To investigate whether NCDN requires the SMN protein for its localisation to vesicles in neural cells, SH-SY5Y cells were transfected with shRNA constructs previously validated to reduce the expression of SMN by an average of 46%, and carrying a GFP marker to unequivocally identify transfected cells (Clelland et al. 2012). Reduction of SMN reduced the number of cytoplasmic foci containing endogenous NCDN (Fig 6A, B). A reduction in the number of nuclear gems (identified with SMN) was also observed, confirming that the shRNA was reducing SMN expression (Fig 6A, C). This, together with data in Figure 5, suggests that NCDN and SMN are mutually dependant for their incorporation into cytoplasmic structures, raising the possibility that the lowered levels of SMN seen in SMA could compromise NCDN function.

**Figure 5:**
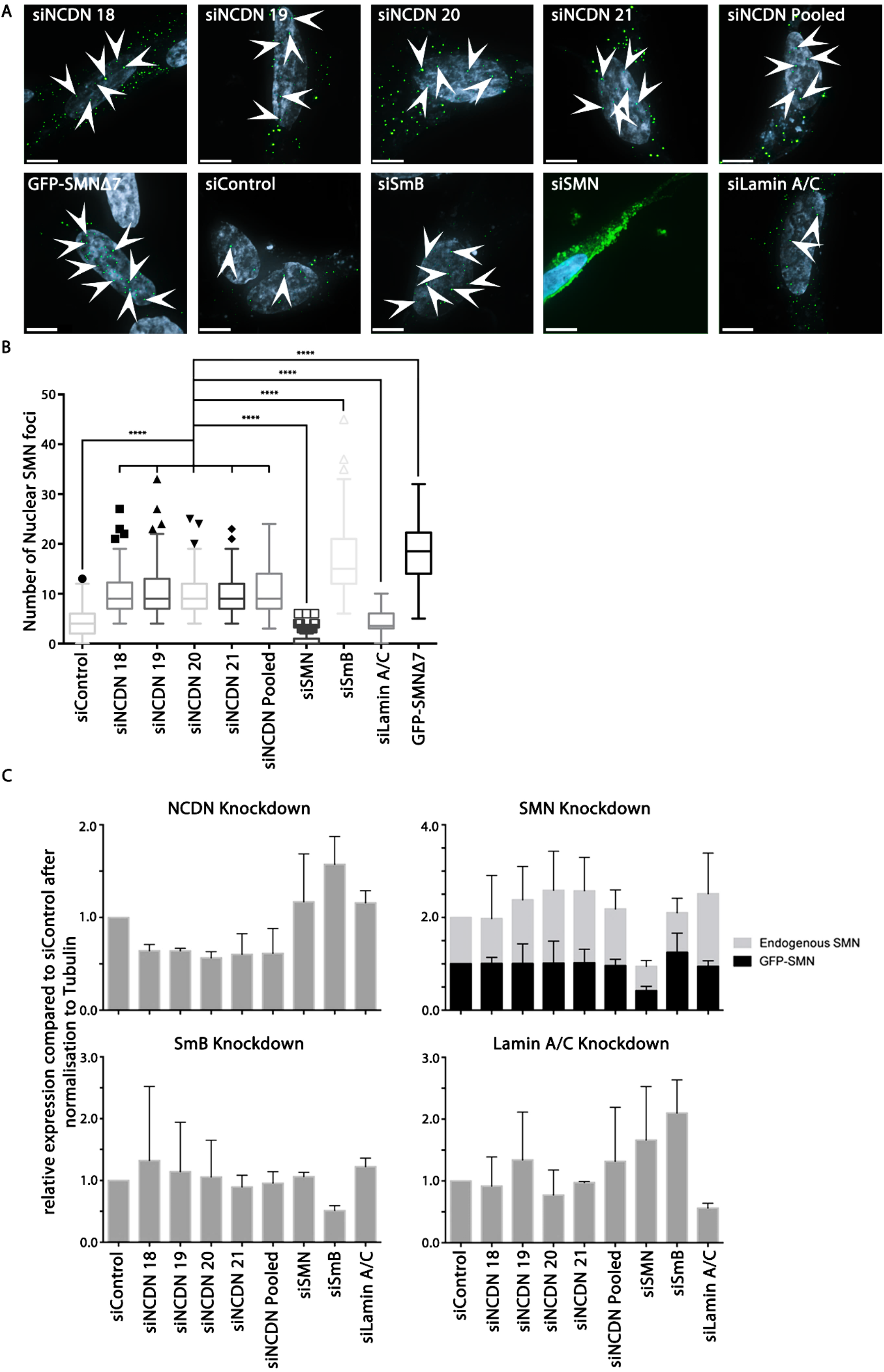
Reduction of endogenous NCDN using siRNA increases localisation of SMN to nuclear gems. *A) Transfection of SH-SY5Y cells constitutively expressing GFP-SMN (Green) with siRNAs shows an increase in the number of nuclear gems (arrows) in cells transfected with siRNAs against NCDN or SmB and a decrease in the number of nuclear gems in cells transfected with siRNAs against SMN in comparison to cells transfected with non-targeting siRNAs (siControl) or siRNAs against lamin A/C as a' targeting' control. Transfection of SH-SY5Y cells with a plasmid to express GFP-SMN∆7 also results in increased numbers of nuclear gems. Cell nuclei are stained with Hoescht 33342 (grey on images). Transfection efficiency with siRNAs was greater than 90%, measured by transfection with siGlo Cyclophillin B (not shown). Bar=7μm. Images are deconvolved z-stacks taken with 0.2μm spacing B) Quantitation of numbers of gems per nucleus shows a significant increase following reduction of NCDN (10.2 (± 4.1) with siNCDN 18, 10.4 (± 4.9)) (Mean ± Standard Deviation) with siNCDN 19, 9.9 (± 4.1) with siNCDN 20, 9.5 (± 3.6) with siNCDN 21 and 10.7 (± 4.6) with siNCDN pooled compared to 4.4 (± 2.5) in cells treated with non-targeting siRNA (siControl) and 4.2 (± 2.3) in cells treated with siRNAs targeting lamon A/C (siLaminA/C). Reduction of SmB also shows an increase in numbers of nuclear gems (to 16.7 (± 6.8) with siSmB), while reduction of SMN leads to a decrease in numbers of nuclear gems (to 0.7 (± 1.4) with siSMN). Expression of GFP-SMN∆7 results in an increase of numbers of nuclear gems to 18.2 (± 5.3). The difference between each siNCDN and controls is statistically significant (AVOVA; P<0.0001, n=150 from 3 replicates). A Tukey post-test identified outliers (individual points marked on graph). C) Immunoblot analysis using antibodies to endogenous NCDN, SMN and SmB shows a reduction in expression of each of 40-60% compared to siControl cells, after signals were normalised to tubulin.*

**Figure 6:**
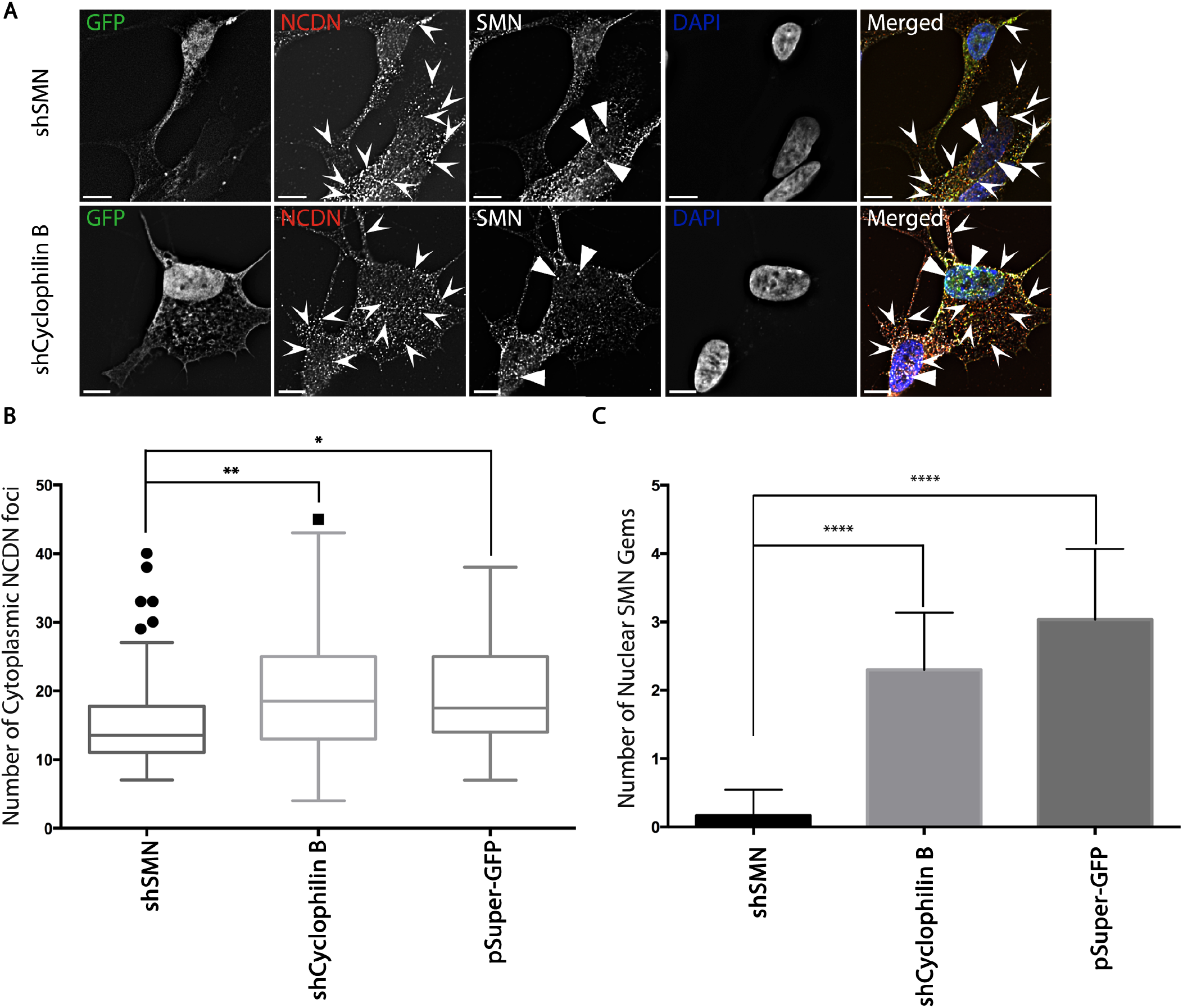
Reduction of endogenous SMN causes a reduction in cytoplasmic NCDN foci in SH-SY5Y cells. *A) SH-SY5Y cells were transfected with plasmids to express shRNAs targeting SMN (shSMN), Cyclophilin B (shCyclophilin) or with the empty pSuper GFP vector (not shown) were fixed after 72 hours, and immunostained for endogenous NCDN and SMN, allowing detection of NCDN foci within the cytoplasm of the cells (identified with Arrowheads), as well as SMN-positive nuclear Gems (identified with Triangles). Bar= 7μm, images are single deconvolved z-sections B) The depletion of SMN results in a reduction in the number of NCDN foci present in the cytoplasm to 15.3 (± 7.2) (Mean ± Standard Deviation) from 20.6 (± 12.0) and 19.5 (± 7.6) compared to cells either transfected with shCyclophilin B or the empty pSuper GFP vector respectively (ANOVA P<0.0005, n=64 from 3 replicates). C) The depletion of SMN causes a reduction of nuclear Gems to 0.17 (± 0.38) from 2.3 (± 0.84) or 3.0 (± 1.0) compared to cells transfected with shCyclophilin B or empty vector respectively (ANOVA P<0.0001, n=30 from 3 replicate).*

### NCDN does not co-purify with splicing snRNPs, suggesting it is not involved in snRNP assembly

To investigate whether the interaction between NCDN and SMN could reflect a previously unidentified role for NCDN in snRNP assembly, splicing snRNPs were affinity purified from whole cells lysates of SH-SY5Y cells constitutively expressing NCDN-GFP using agarose beads coupled to antibodies against the characteristic tri-methyl guanosine Cap of snRNAs (TMG beads, Millipore) (Fig 7). Endogenous Sm proteins showed strong enrichment in the affinity purified snRNP samples, as expected for a core snRNP protein. Endogenous SMN was also co-enriched, demonstrating that the experimental conditions were suitable to identify proteins important for snRNP assembly. NCDN-GFP did not co-purify with splicing snRNPs, suggesting that NCDN is not involved in snRNP assembly or processing. This raises the intriguing possibility that the interaction between SMN and NCDN reflects a novel, snRNP-independent role for SMN.

**Figure 7:**
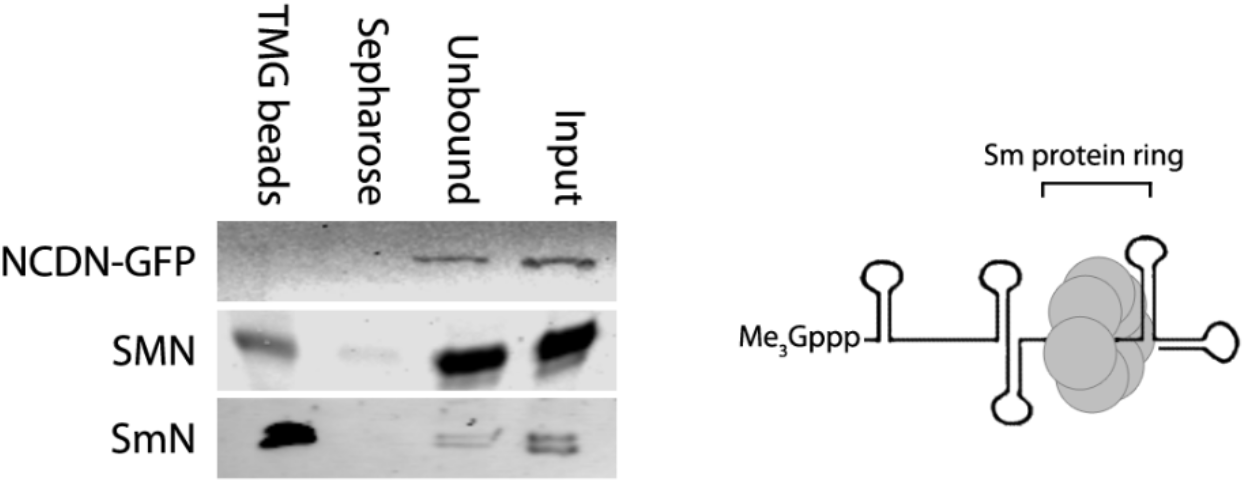
NCDN-GFP does not co-purify with snRNPs in SH-SY5Y cells constitutively expressing NCDN-GFP. *Incubation of whole-cell lysate from an SH-SY5Y cell line constitutively expressing NCDN-GFP with agarose beads conjugated to antibodies against the tri-methyl guanosine cap (Me3Gppp) of snRNAs (TMG beads) affinity purifies snRNPs as evidenced by the enrichment of the core snRNP protein SmN (detected with anti-SmN, bottom row). The enriched snRNP fraction also contains SMN, which is essential for snRNP assembly. NCDN-GFP,however, does not co-enrich with snRNPs. Also shown is the core structure of mature snRNPs consisting of the heptameric Sm protein ring bound at the Sm binding site of snRNA, as well as the characteristic tri-methyl guanosine Cap of snRNAs (Me3Gppp) at the 5' end.*

### SMN interacts with Rab5 in SH-SY5Y cells and co-localises with a sub-set of Rab5 vesicles

Recent studies have found that SMA may cause endocytic defects, especially in synaptic vesicle recycling in animal models (Dimitriadi et al. 2016). Several SMA-protective disease modifier genes, such as Plastin 3, Coronin 1C, and Neurocalcin Delta, are also associated with endocytosis (Oprea et al. 2008, Hosseinibarkooie et al. 2016, Riessland et al. 2017). However, other endocytic structures within the cell have not been investigated. Rab5 is a marker of early endosomes and endocytic vesicles, as well as being a regulator of these trafficking pathways (Bucci et al. 1992). NCDN and Rab5 have previously been shown to interact, while both Rab5 and NCDN both have roles in dendrite morphogenesis and cell polarity (Satoh et al. 2008, Oku et al. 2013, Guo et al. 2016).

As we had previously shown that SMN and NCDN co-localise in vesicles, we hypothesised that some of the SMN-rich vesicles could be Rab5 vesicles. SH-SY5Y cells were co-transfected with plasmids to express mRFP-Rab5 (Vonderheit, Helenius 2005) together with GFP-SMN, NCDN-GFP or YFP. mRFP-Rab5 was affinity-purified from whole cell lysates from each co-transfection using RFP-TRAP (Chromotek). Subsequent immunoblotting revealed co-purification of GFP-SMN and NCDN-GFP, but not YFP alone, with mRFP-Rab5 (Fig 8A). Furthermore, endogenous SMN also co-purified with mRFP-Rab5. In parallel experiments, co-localisation of mRFP-Rab5 with GFP-SMN and NCDN was investigated (Fig 8B, C). In accordance with previous publications, Rab5 showed partial co-localisation with NCDN-GFP in cytoplasmic structures (arrows in Fig 8C) (Oku et al. 2013). GFP-SMN showed a similar degree of co-localisation with mRFP-Rab5, also in cytoplasmic structures, while there was minimal co-localisation between YFP and mRFP-Rab5. Taken together with the data in Figure 7 above this suggests that NCDN and SMN co-localise in the context of Rab5 vesicles, independently of snRNP assembly.

**Figure 8:**
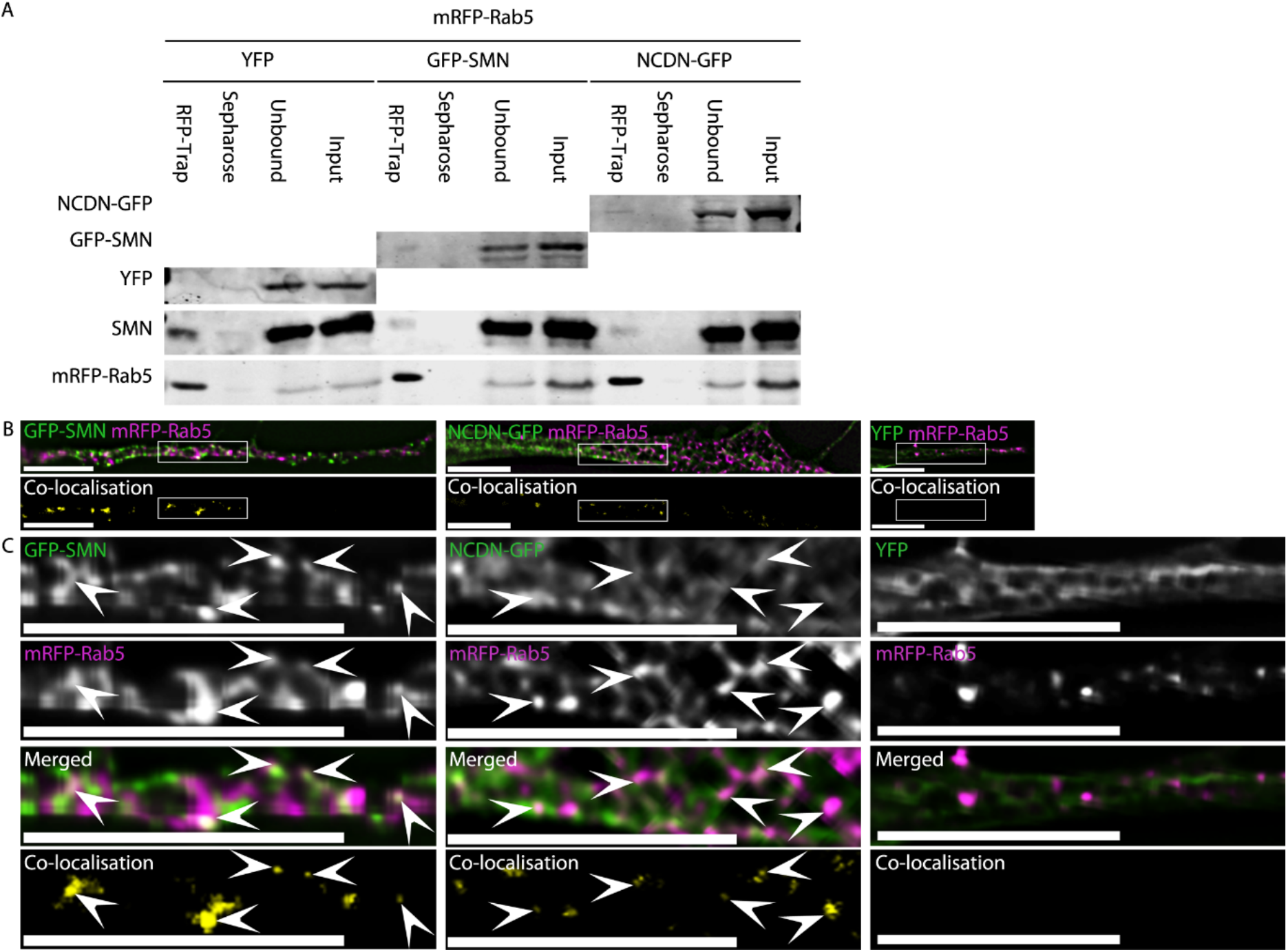
NCDN and SMN interact with Rab5 and co-localise with a subset of Rab5 vesicles within neurites of SH-SY5Y cells. M*A) Affinity isolation of mRFP-Rab5 using RFP-Trap from cells co-transfected with plasmids to express mRFP-Rab5 together with NCDN-GFP, GFP-SMN or YFP alone co-enriches both NCDN-GFP (top row, detected with anti-GFP, band is present in RFP-Trap lane but not sepharose beads lane) and SMN-GFP (second row, detected with anti-GFP, band is present in RFP-Trap lane but not sepharose beads lane), but not YFP (third row, no band detected in RFP-Trap lane). Endogenous SMN (fourth row, detected with mouse anti-SMN) co-enriches with mRFP-Rab5 in all three samples. Detection of mRFP-Rab5 (bottom row, detected with anti-RFP) confirms robust enrichment of mRFP-Rab5 in all three samples. B) Both GFP-SMN and NCDN-GFP partially co-localise with mRFP-Rab5 in a subset of mRFP-Rab5 containing vesicles in co-transfected SH-SY5Y cells (white signal in overlaid images, top row; yellow signal in co-localisation images, bottom row). C) Enlargement of the boxed areas in B confirm that the co-localisation between SMN/NCDN and Rab5 occurs in punctate structures. Arrowheads identify areas of co-localisation. Co-localisation images were generated by Volocity, using automatic thresholds on non-deconvolved z-sections before excluding values below 0.05. Images (excluding the co-localisation images) are single deconvolved z-sections. Bar=7μm.*

## Discussion

The genetic cause of SMA has been known since 1995 (Lefebvre et al. 1995), but there is still little available in the way of treatment. Spinraza/Nusinersin is now available to treat SMA by correcting the defective splicing of the SMN2 transcript to promote production of full-length SMN protein. However, this is not a complete cure and requires regular maintenance doses through intrathecal injection. Additionally, little has been done to investigate potential symptoms that could arise later in life or in other tissues and organs in patients treated with Spinraza. Additional treatment options for SMA are still needed, for use in addition to Spinraza, or in place of it for patients for whom it is not suitable, including those with more rare forms of SMA in which SMN is not mutated.

A significant reason for the lack of treatment options for SMA is uncertainty about the cellular roles of SMN, which appear to be numerous. In particular, it is not clear why motor neurones are so exquisitely sensitive to reduced levels of SMN when the key roles of the protein appear to be in pathways required in all cell types. By comparing the interactomes of two very similar members of Sm protein family, SmB and the neural-specific SmN, we have uncovered an interaction between SMN and the essential neural protein NCDN, which may open novel avenues for therapy development.

### The neural-specific Sm protein, SmN, behaves similarly to SmB, but shows subtle differences at the interactome level that may indicate alternative roles

Differences between members of the Sm protein family have not been systematically investigated, although non-splicing roles have been proposed for SmB and SmD3 in mRNA localisation and for SmD1 in miRNA biogenesis in Drosophila (Gonsalvez et al. 2010, Xiong et al. 2015). As SmB and SmN are thought to perform the same primary function in snRNPs (Huntriss et al. 1993), it is currently unknown why SmN is expressed in neural tissues as well as, or instead of, SmB. Current research has suggested that the expression of SmN may cause tissue specific alternative splicing of pre-mRNA transcripts (Lee et al. 2014). However, an alternative, but complimentary hypothesis is that SmN may be adapted for secondary, neural specific roles. We demonstrate here that SmN localises identically to SmB during snRNP maturation and at steady-state where both localise to vesicles containing SMN in the cytoplasm and neurites of SH-SY5Y cells in addition to their canonical localisation to nuclear CBs and speckles. Our parallel proteomic study used SH-SY5Y neural cell lines constitutively expressing YFP-SmN and YFP-SmB to investigate difference between the interactomes of these two, very similar, proteins. YFP-SmN has a proline rich C-terminal tail that YFP-SmB lacks, although a similar sequence is present in SmB' (Mourão et al. 2016), an alternatively-spliced product of the SNRPB gene, which encodes SmB. Several proteins were identified in the SmN interactome but not the SmB interactome such as nuclear receptor co-activator 6 interacting protein (UniProt Q96RS0), and 7SK snRNA methylphosphate capping enzyme (UniProt Q7L2J0), both of which are associated with snRNA capping (Jeronimo et al. 2007, Hausmann et al. 2008). Additionally, RNA-binding protein 40 (UniProt Q96LT9) was uniquely identified within the SmN interactome, and is involved in the minor spliceosome (Benecke et al. 2005). This suggests that some of these proteins may interact preferentially with SmN, perhaps mediated by amino acid changes within the proline-rich tail. However, further validation and additional experimentation would be required to confirm these differences in interactome between SmN and SmB and to investigate specific functions for the distinct Sm protein family members.

### NCDN interacts with SMN, SmB and SmN and co-localises with them in vesicles, suggesting a novel cellular role for SMN

Previous research into neural-specific functions for SMN has identified several new protein-protein interactions involving SMN. These novel SMN partners have, in the main, been RNA binding proteins (Rossoll et al. 2003b, Fallini et al. 2011, Akten et al. 2011, Fallini et al. 2014). There is growing appreciation that SMN-mediated transport may be of particular importance in neural cells and involve COP1-type vesicles transported by Dynein and containing SmB (Peter et al. 2011, Custer et al. 2013, Prescott et al. 2014, Li et al. 2015). The nature and content of these vesicles is not clear but they are likely to be of significance for the cell-type bias of SMA symptoms, as they are present predominantly in neural cells (Rossoll et al. 2003a, Zhang et al. 2003, Zhang et al. 2006a, Todd et al. 2010a, Todd et al. 2010b, Akten et al. 2011, Fallini et al. 2011, Peter et al. 2011, Custer et al. 2013, Fallini et al. 2014, Prescott et al. 2014, Li et al. 2015, Fallini et al. 2016). The SmN/SmB interactome screen presented here suggests a large number of non-snRNP proteins as potential cellular partners for the Sm proteins.

We chose to investigate the neural protein NCDN further, as it has characteristics that may be of relevance for SMA. NCDN is predominantly expressed in neural tissue, and little is known about its structure or function, as it shares little sequence homology with other eukaryotic proteins (Shinozaki et al, 1997). Though characterised relatively poorly, it is associated with dendrite morphogenesis and localises to Rab5 vesicles involved in the maintenance of cell polarity (Oku et al. 2013, Guo et al. 2016), NCDN has also been demonstrated to regulate localisation of signalling proteins such as P-Rex 1 (Pan et al. 2016), suggesting that it, in common with SMN, has a role in intra-cellular trafficking. These neural-specific and trafficking roles suggest that further analysis of the interaction between NCDN and the Sm proteins may help to better understand the molecular mechanisms of pathogenesis in SMA.

Reciprocal affinity-purification of GFP-tagged and endogenous NCDN and Sm proteins (Fig 3A, B) validated the interaction detected in the interactome analysis. Although originally identified as a protein interacting with SmN but not SmB, further investigation indicates that NCDN is, in fact, capable of interacting with both of these Sm proteins. Of much greater interest, however, is the interaction documented between NCDN and SMN, which appears more robust than that between NCDN and the Sm proteins (Fig 3B, E). Furthermore, NCDN is excluded from the cell nucleus and localises with SMN and the Sm proteins in mobile vesicles in the neurites of SH-SY5Y cells (Fig 3D, F) suggesting that it shares cytoplasmic, rather than nuclear, roles with SMN. The truncated protein SMN∆7, which can't fully substitute for FL-SMN, is largely restricted to the nucleus (Renvoisé et al. 2006, Sleigh et al. 2011). While it can't yet be ruled out that SMN∆7 lacks the capability to substitute for FL-SMN in nuclear roles, this suggests that cytoplasmic roles of SMN are key to SMA pathology. NCDN was not co-purified with splicing snRNPs, under conditions that showed a clear enrichment of SMN in the snRNP fraction. This suggests that the interaction between NCDN, SMN and the Sm proteins is not related to snRNP assembly.

### Potential consequences of NCDN mis-localisation associated with SMN reduction

We have identified co-localisation of both SMN and NCDN with a sub-set of Rab5 vesicles. Since NCDN is also found in a sub-set of SMN-positive cytoplasmic structures, it is highly likely that these are Rab5 vesicles. It is possible that the protein-protein interactions between NCDN and SMN occur elsewhere in the cytoplasm as both proteins also show a diffuse cytosolic pool, but the decrease seen in cytoplasmic structures containing NCDN following SMN depletion suggest that cellular pathways requiring NCDN-containing vesicles may be compromised in SMA.

At present, the precise roles of NCDN are not fully understood, although it has been implicated in dendrite morphogenesis, neural outgrowth, synaptic plasticity regulation, and moderation of signalling pathways in neural cells (Shinozaki et al. 1997, Shinozaki et al. 1999, Ohoka et al, 2001, Dateki et al. 2005, Francke et al. 2006, Wang et al. 2009, Ward et al. 2009, Oku et al. 2013, Wang et al. 2013, Matosin et al. 2015, Pan et al. 2016). NCDN has also previously been shown to localise to Rab5 vesicles within dendrites (Oku et al. 2013). These dendritic Rab5 vesicles have been found have an important role in dendrite morphogenesis and somatodendritic polarity (Satoh et al. 2008, Guo et al. 2016). As we have now demonstrated that SMN localises to a sub-set of Rab5 vesicles, likely in association with NCDN, SMN may also be implicated in cell polarity, with an insufficiency of SMN causing problems with both establishment and maintenance of polarity. These would be particularly vital in such elongated cells as motor neurons and may be mediated through trafficking of mRNAs or proteins. Further work will be required to investigate defects in cell polarity as a pathogenic mechanism in SMA and their possible link to NCDN.

### NCDN as a novel therapeutic target in SMA

SMN has now been linked to several functions other than its canonical role in snRNP assembly. While reduction in the cell's capacity for snRNP assembly caused by lowered SMN may cause splicing defects, the key transcripts preferentially affecting motor neurones are still to be identified. SMN has an established role in the trafficking of mature mRNAs destined for localised translation. The nature of the structures involved in this role is not completely clear, however, with different authors describing the structures as vesicular or granular. Reduction of SMN has also been linked with endosomal defects, suggestive of the importance of SMN for vesicular transport. Here we provide further evidence for the presence of SMN in, or associated with, vesicles and identify the essential neural protein, NCDN as a potential target for therapy development in SMA. Further work will be required to establish which roles of NCDN also involve SMN, however the co-dependence of SMN and NCDN in cytoplasmic vesicles suggests that depletion of SMN, as seen in the majority of SMA patients, may affect NCDN localisation and/or function. Manipulation of the expression or function of NCDN may, therefore, have potential as an SMN-independent target for the development of novel therapies for SMA.

## Materials and Methods

### Plasmid constructs

pEGFP-SMN, pEYFP-SmB and mCherry-SmB have been described previously (Sleeman, Lamond 1999, Sleeman et al. 2001, Clelland et al. 2009). pEYFP-SmN and pmCherry-SmN were generated by PCR amplification and sub-cloning cDNA of human SmN from SH-SY5Y cells into pEYFP-C1 and pmCherry-C1 respectively, using SNRPNEcoRI forward primer: TAGAATTCCATGACTGTTGGCAAGAGTAGC, and SNRPNBamHI reverse primer: TAGGATCCCTGAGATGGATCAACAGTATG. pmCherry-SMN was generated by PCR amplification subcloning the sequence from the pEGFP-SMN plasmid into pmCherry-C1 using an SMNEcoRI Forward primer: GCGGAATTCTATGGCGATGAGC and SMNBamHI Reverse Primer: GCAGGATCCTTAATTTAAGGAATGTGA. To generate pEGFP-NCDN, NCDN cDNA from SH-SY5Y cells was PCR amplified and sub-cloned into a pEGFP-N3 plasmid using NCDNEcoRI forward primer: GCGGAATTCATGGCCTCGGATTGCG and NCDNSalI reverse primer: GCTGCTGACGGGCTCTGACAGGC. All cDNAs were amplified using GoTaq G2 (Promega) and the PCR products restriction digested using EcoRI and either BamHI or SalI (Promega), before ligation with T4 DNA ligase (Thermo Scientific). mRFP-Rab5 was a gift from Ari Helenius (Vonderheit, Helenius 2005).

### Cell lines and cell culture

SH-SY5Y cells were from ATCC. Cells were cultured in DMEM with 10% FBS at 37°C, 5% CO2. Transfections were carried out using Effectene (Qiagen) according to the manufacturer's instructions. Stable SH-SY5Y cell lines expressing mCherry–SmB and GFP–SMN have been described previously (Clelland et al. 2009, Prescott et al. 2014). SH-SY5Y cell lines stably expressing YFP-SmN, YFP-SmB, YFP, mCherry-SmN and NCDN-GFP were derived by clonal isolation following selection with 200 μg/ml G418 (Roche) following transfection.

### Immunostaining, microscopy and image analysis

Cell fixing and immunostaining were both carried out as described previously (Sleeman et al. 2003). Live cell and fixed cell microscopy and image processing were carried out as described previously (Prescott et al. 2014). BODIPY-493 (Life Technologies) was added to culture medium at 2 μg/ml overnight. Antibodies used for immunostaining were Mouse monoclonal Y12 anti-Smith (SmB) (Abcam ab3138, 1:20), Rabbit polyclonal 204-10 (anti-Coilin) (a gift from A. I. Lamond (Bohmann et al. 1995), 1:500), Mouse monoclonal anti-SMN (BD Transduction 610646, 1:50) and Rabbit polyclonal anti-NCDN (Proteintech 13187-1-AP, 1:50). Overlays of images were made using Adobe Photoshop CS5 (Adobe). Co-localisation images were generated using Volocity 6.3 (PerkinElmer), using automatic thresholds on non-deconvolved images. Co-localisation values of 0.05 or less were excluded as this was the maximum co-localisation value observed between mCherry signal and YFP signal in neurites expressing YFP as a control together with mCherry-tagged proteins of interest. Deconvolution was also performed using Volocity 6.3, with between 15 and 25 iterations of deconvolution.

### Statistical Analysis, and generation of graphs

Data was processed using Microsoft Excel (Microsoft) to produce ratios, proportions and percentages. Bar charts, Box and Whisker plots were then generated using Prism 6 (GraphPad) from the processed data. Statistical analysis was also performed using Prism 6, with multiple comparisons to determine statistical difference between specific sets of data. Tukey post-tests were used to identify outliers in Anova statistical analysis.

### Preparation of cell lysates and immunoblotting

Cells were grown in 10cm diameter dishes, before being detached with trypsin and collected by centrifugation at 180 RCF for 5 minutes. The cell pellet was washed 3 times in PBS before lysis in 100μl of ice cold lysis buffer per dish (50 mM Tris-HCl pH 7.5; 0.5 M NaCl; 1% (v/v) Nonidet P-40; 1% (w/v) sodium deoxycholate; 0.1% (w/v) SDS; 2 mM EDTA plus cOmplete mini EDTA-free protease inhibitor cocktail (Roche, one tablet per 10 ml)), followed by homogenisation by sonication. Isolation of YFP/GFP and mCherry/mRFP-tagged proteins was carried out as described previously with GFP - or RFP-Trap (Chromotek) (Prescott et al. 2014). Lysates were electrophoresed on a 10% SDS-polyacrylamide gel and transferred to nitro-cellulose (Hybond-C+ or Protran premium 0.2μm, both GE Healthcare) membranes for immunoblotting. Antibodies used were rat monoclonal anti-RFP (Chromotek 5F8, 1∶500); goat polyclonal anti-γCOP (Santa Cruz sc-14167, 1∶250), rabbit polyclonal anti-GFP (Abcam ab290, 1:2000), rabbit polyclonal anti-SNRPN (SmN) (Proteintech 11070-1-AP, 1:800), mouse monoclonal Y12 anti-Smith (SmB) (Abcam ab3138, 1:100), rabbit polyclonal anti-SMN (Santa Cruz sc-15320, 1:500), mouse monoclonal anti-SMN (BD Transduction labs 610646, 1:500), rabbit polyclonal anti-COPB1 (CUSAB CSB-PA005783LA01HU, 1:500), mouse monoclonal anti-Lamin A/C (Santa Cruz sc-7292, 1:500), rabbit polyclonal anti-NCDN (Proteintech 13187-1-AP, 1:500), mouse Monoclonal anti-tubulin (Sigma Aldrich, 1:500) and rabbit polyclonal anti-Histone H3 (Proteintech 17168-1-AP, 1:300). Secondary antibodies were goat anti-rabbit Dylight 700 (Thermo Scientific 35569, 1:15,000) or goat anti-mouse Dylight 800 (Thermo Scientific SA5-10176, 1:15,000). Alternatively, goat anti-mouse IRDye 800CW (Li-Cor 925-32210, 1:25,000) and goat anti-rabbit IRDye 680RD (Li-Cor 925-68071, 1:25,000) were used. Goat anti-Rat Dylight 800 (Thermo Scientific, SA5-10024) antibody was used to visualise the rat monoclonal anti-RFP antibody at a concentration of 1:15,000. Donkey anti-goat IRDye 800CW (Licor, 925-32214) was used at a concentration of 1:25,000 to detect goat polyclonal anti-γCOP. Donkey anti-rabbit conjugated to horseradish peroxidase (HRP) (Pierce, 31460, 1:15,000) was used to identify endogenous NCDN in figure 3D. Detection of antibodies conjugated to fluorophores was carried out with an Odessey CLx using Image Studio (both Li-cor). Band quantification was also performed using Image Studio. Detection of antibodies conjugated to peroxidase was performed using ECL Western Blotting Substrate (Pierce) and developed with Hyperfilm (Amersham), using a Kodak X-OMAT 1000 developer, after 30-45 minutes exposure.

### Immunoprecipitation of intact snRNPs

To immunoprecipitate intact snRNPS, whole cell lysates were incubated with anti-2,2,7-trimethylguanosine conjugated to agarose beads (Millipore NA02A), with Sepharose 4B beads (Sigma Aldrich) as a control. 40 ng of pre-cleared lysate and unbound protein were separated by SDS-PAGE alongside material precipitated with Sepharose control beads and TMG antibody beads. Subsequent detection was carried out using rabbit anti-GFP (1:2000, Abcam), rat mAb anti-RFP (1:500, Chromotek), mouse monoclonal anti-SMN (1:500, BD Transduction labs) and rabbit polyclonal anti-SNRPN (1:800, Proteintech).

### Preparations and analysis of Mass Spectrometry samples

SH-SY5Y cells constitutively expressing either YFP, YFP-SmN or YFP-SmB were lysed in co immunoprecipitation buffer (10mM Tris pH7.5, 150mM NaCl, 0.5mM EDTA, 0.5% NP40, 1 cOmplete EDTA-free protease inhibitor tablet (Roche) per 10ml), followed by affinity purification of the tagged proteins with GFP-Trap as above. 5μl of the affinity isolated material, alongside precleared lysate and unbound lysate was transferred to nitrocellulose membrane (as above) and immunodetected using Rabbit anti-GFP (Abcam) to confirm efficient immunoprecipitation. Samples were then electrophoresed on a NuPAGE 4-12% Bis-Tris Acrylamide gel (Novex NP0321), Coomassie stained using SimplyBlue SafeStain (Invitrogen), gel chunks excised and analysed by the Mass Spectrometry and Proteomics Facility at the University of St Andrews.

The gel chunks were cut into 1 mm cubes. These were then subjected to in-gel digestion, using a ProGest Investigator in-gel digestion robot (Digilab) using standard protocols (Shevchenko et al. 1996). Briefly, the gel cubes were destained by washing with MeCN and subjected to reduction with DTT and alkylation with IAA before digestion overnight with trypsin at 37°C. The peptides were extracted with 10% formic acid, and the volume reduced to ~20ul by concentration in a speedvac (ThermoSavant).

The peptides were then injected onto an Acclaim PepMap 100 C18 trap and an Acclaim PepMap RSLC C18 column (ThermoFisher Scientific), using a nanoLC Ultra 2D plus loading pump and nanoLC as-2 autosampler (Eksigent). The peptides were eluted with a gradient of increasing acetonitrile, containing 0.1 % formic acid (2-20% acetonitrile in 90 min, 20-40% in a further 30 min, followed by 98% acetonitrile to clean the column, before re-equilibration to 2% acetonitrile). The eluate was sprayed directly into a TripleTOF 5600 electrospray tandem mass spectrometer (Sciex, Foster City, CA) and analysed in Information Dependent Acquisition (IDA) mode, performing 250 msec of MS followed by 100 msec MS/MS analyses on the 20 most intense peaks seen by MS. The MS/MS data file generated via the ‘Create mgf file' script in PeakView (Sciex) was analysed using the Mascot search algorithm (Matrix Science), against the NCBInr database (Oct 2014) restricting the search to Homo sapiens (284,317 sequences), trypsin as the cleavage enzyme and carbamidomethyl as a fixed modification of cysteines and methionine oxidation as a variable modification. The peptide mass tolerance was set to ± 0.05 Da and the MSMS mass tolerance to ± 0.1 Da.

Scaffold viewer (version Scaffold_4.5.1, Proteome Software) was used to validate MS/MS based peptide and protein identifications. Peptide identifications were accepted if they could be established at greater than 95.0% probability by the Peptide Prophet algorithm (Keller et al. 2002). Protein identifications were accepted if they could be established at greater than 99.0% probability and contained at least 2 identified peptides. Protein probabilities were assigned by the Protein Prophet algorithm (Nesvizhskii et al. 2003). Proteins that contained similar peptides and could not be differentiated based on MS/MS analysis alone were grouped to satisfy the principles of parsimony.

Identified proteins affinity purified alongside YFP-SmB or YFP-SmN were discounted if they were additionally identified as being affinity purified with YFP, or if they were present within the Sepharose bead proteome (Trinkle-Mulcahy et al. 2008).

### RNAi assays

Reduction of protein expression using siRNA was achieved by transfecting the appropriate cell lines with siRNAs (Dharmacon) using Viromer Green (Lipocalyx GmbH) according to the manufacturer's instructions. Cells were lysed for assay by immunoblotting, or fixed with paraformaldehyde for fluorescence microscopy, 48 hours after transfection. Sequences used were SMN: CAGUGGAAAGUUGGGGACA; SmB, a mixture of CCCACAAGGAAGAGGUACU, GCAUAUUGAUUACAGGAUG, CCGUAAGGCUGUACAUAGU, CAAUGACAGUAGAGGGACC; NCDN, individually and a mixture of NCDN 18 GUUCAUUGGUGACGAGAAA, NCDN 19 AGACCUCAUCCUUGCGUAA, NCDN 20 AGGCCAAGAAUGACAGCGA, NCDN 21 GGCCAUUGAUAUCGCAGUU; negative control (siControl) targeting luciferase, UAAGGCUAUGAAGAGAUAC; positive control targeting Lamin A/C, GGUGGUGACGAUCUGGGCU; SiGlo Cyclophillin B to determine transfection efficiency, GGAAAGACUGUUCCAAAAA. Lysates were electrophoresed on an SDS-PAGE gel, transferred to nitrocellulose membrane, and immunodetected with antibodies to the above proteins. Band signal intensity determined with ImageStudio (Li-Cor), and the values were normalised to Tubulin.Reduction of protein expression using shRNA was achieved by transfecting SH-SH5Y cell lines with pSUPER-GFP.Neo plasmids (Oligoengine) expressing shRNA to SMN and Cyclophilin B, which have been described previously (Clelland et al. 2012) using Effectene (QIAGEN).

### Fractionation

Cells were pelleted from the appropriate cell line, and incubated in Buffer A (10mM HEPES pH7.9, 1.5mM MgCl_2_, 10mM KCl, 0.5mM DTT, 1 cOmplete EDTA-free protease inhibitor tablet per 10ml) for 5 minutes, before being Dounce homogenised 25 times using the tight pestle to disrupt the plasma membrane. This was then centrifuged at 300 RCF for 5 minutes to pellet the nuclei. The supernatant was removed, recentrifuged at 300 RCF to further remove nuclei, before the supernatant was centrifuged at 16,100 RCF for 30 minutes using a refrigerated 5415R Centrifuge (Eppendorf). The nuclei were resuspended in Buffer S1 (250mM Sucrose, 10mM MgCl_2_), before this was layered over with Buffer S3 (880mM Sucrose, 0.5mM MgCl_2_). The nuclear pellet was then centrifuged at 2800 RCF for 10 minutes to wash and pellet the nuclei. The supernatant from the 16,100 RCF centrifugation was further centrifuged at 100,000 RCF using an Optima Max-XP ultracentrifuge with a TLA-110 Rotor (Beckman-Coulter) for 60 minutes. The supernatant was removed and kept. The 16,100 and 100,000 RCF pellets were washed in Buffer A and centrifuged at 16,100 RCF for 30 minutes or 100,000 RCF for 1 hour respectively. Each pellet was then resuspended in lysis buffer (see above). To confirm efficient separation of cytoplasmic fractions from the nuclear fractions, Mouse anti-tubulin (Sigma Aldrich, 1:500) and Rabbit polyclonal anti Histone H3 (Proteintech, 1:300) antibodies were used.

## Acknowledgements

The authors thank Prof. Ari Helenius (ETH Zurich) for the mRFP-Rab5 construct, Prof Gary Bassell, Emory University for the pGFP-SMN∆7 construct and Prof. Angus Lamond (University of Dundee) for anti-coilin antibodies.

## Author contributions

LWT designed experiments, generated reagents, produced and analysed data, generated the figures and helped to prepare and edit the manuscript. KM acquired and analysed data. SS and CB processed the MS samples, produced the MS/MS data set and assisted with analysis. JES came up with the original concept, designed experiments, analysed data and prepared and edited the manuscript.

---

**Figure S1:**
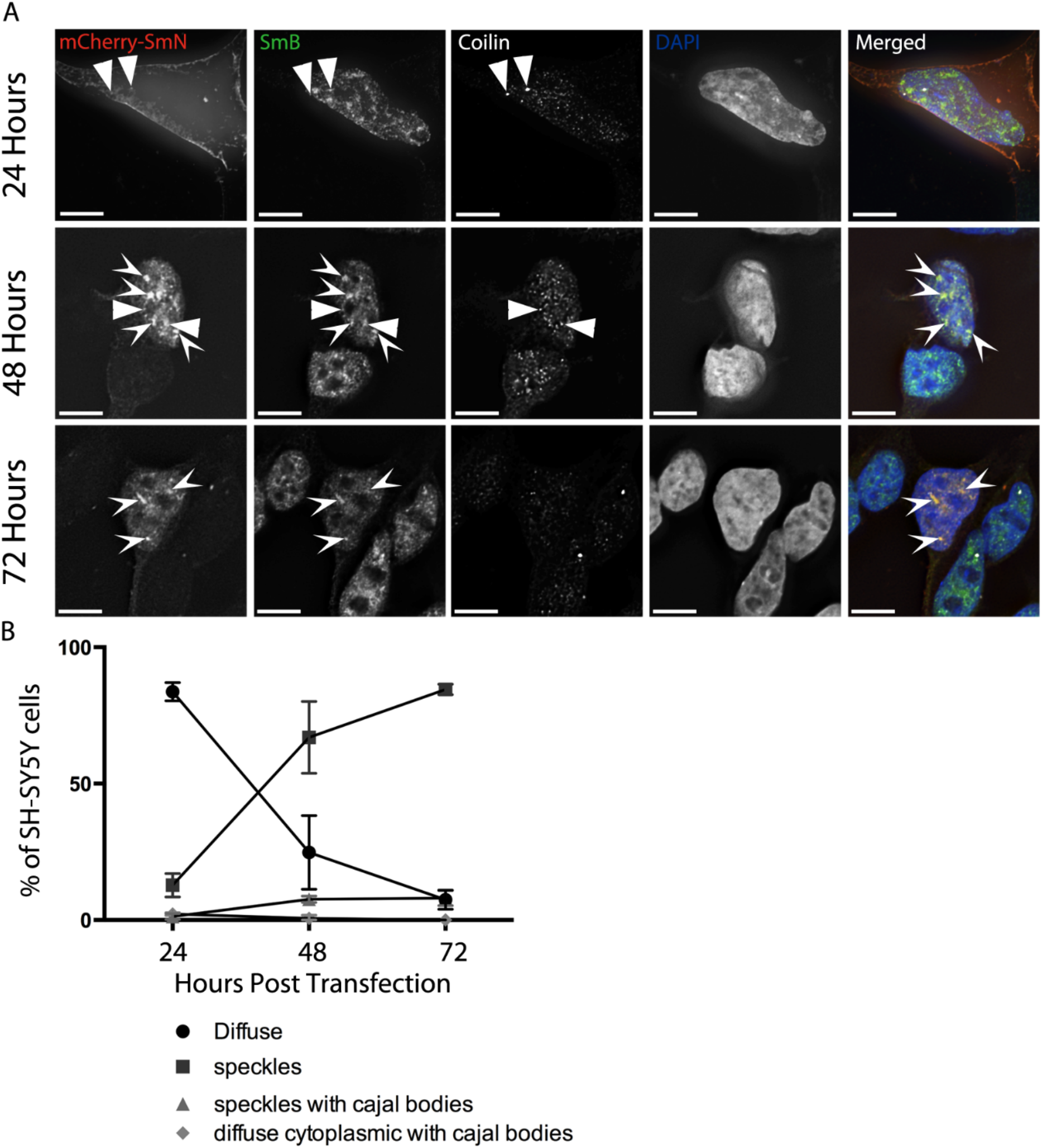
*mCherry-SmN exhibits similar behaviour to both YFP-SmN and SmB in SH-SY5Y cells. A) SH-SY5Y cells transiently expressing mCherry-SmN and fixed after 24, 48 and 72 hours show variations in distribution of the mCherry-SmN with time. Immunostaining with Y12 (green on overlay) and anti-coilin (white on overlay) shows splicing speckles (arrowheads) and Cajal Bodies (CBs, triangles) respectively. Images are deconvolved z-stacks with 0.2 μm spacing. Bar=7μm. B) mCherry-SmN initially localises diffusely in the cytoplasm, before localising to speckles at the 48 and 72 hour time-points. 3 independent experiments, n=100 cells per experiment. Data shown is mean ± SD.*

**Table S2:** *A selected dataset from the interactome analysis of SmN and SmB confirms efficient identification of known Sm protein interactors. All other core Sm family proteins were identified in the interactome analysis, as well as SMN, all Gemin components of the SMN complex and several LSm proteins, including LSm11 (found only in the U7 snRNP). Additionally, several members of the methylosome, where SmB post-translational modifications occur, were identified including Protein arginine N-methyltransferase 5 (PRMT5). Several previously identified SmB interactors (CD2 antigen cytoplasmic tail binding protein-2, PERQ2, WW domain-binding protein 4 and Formin binding protein 4) (Bedford et al. 1998, Bedford et al. 2000, Kofler et al. 2004, Kofler et al. 2005) were identified to interact with at least one of the proteins. * denotes that spectra from SmN were pooled with those from SmB by Scaffold due to sequence similarity in the majority of the proteins. Neurochondrin, and other proteins discussed are also shown here. Values are number of unique peptides identified.*

| Identified protein | Accession number | Molecular Weight | SmN | SmB | YFP |
| --- | --- | --- | --- | --- | --- |
| small nuclear ribonucleoprotein polypeptide B * [Homo sapiens] | gi\|53690154 | 24 kDa | 13 | 11 | 2 |
| small nuclear ribonucleoprotein Sm D2 isoform 1 [Homo sapiens] | gi\|4759158 | 14 kDa | 10 | 8 | 4 |
| small nuclear ribonucleoprotein Sm D1 [Homo sapiens] | gi\|5902102 | 13 kDa | 5 | 5 | 4 |
| small nuclear ribonucleoprotein E [Homo sapiens] | gi\|4507129 | 11 kDa | 5 | 4 | 2 |
| small nuclear ribonucleoprotein F [Homo sapiens] | gi\|4507131 | 10 kDa | 5 | 3 | 0 |
| small nuclear ribonucleoprotein Sm D3 [Homo sapiens] | gi\|4759160 | 14 kDa | 4 | 4 | 3 |
| small nuclear ribonucleoprotein G [Homo sapiens] | gi\|4507133 | 8 kDa | 4 | 3 | 0 |
| survival motor neuron protein isoform a [Homo sapiens] | gi\|13259527 | 30 kDa | 8 | 7 | 0 |
| GEMIN5 protein [Homo sapiens] | gi\|219517971 | 168 kDa | 22 | 25 | 0 |
| HC56 (Gemin 4) [Homo sapiens] | gi\|10945430 | 119 kDa | 18 | 11 | 0 |
| DEAD box RNA helicase Gemin3 [Homo sapiens] | gi\|6503002 | 92 kDa | 12 | 18 | 0 |
| gem-associated protein 2 isoform alpha [Homo sapiens] | gi\|4506961 | 32 kDa | 6 | 4 | 0 |
| gem-associated protein 8 [Homo sapiens] | gi\|8923481 | 29 kDa | 6 | 2 | 0 |
| Gem (nuclear organelle) associated protein 6 [Homo sapiens] | gi\|17390437 | 19 kDa | 3 | 3 | 0 |
| gem-associated protein 7 [Homo sapiens] | gi\|13376001 | 15 kDa | 3 | 2 | 0 |
| WD-40 repeat protein (Unrip) [Homo sapiens] | gi\|4519417 | 38 kDa | 16 | 16 | 0 |
| U6 snRNA-associated Sm-like protein LSm4 isoform 1 [Homo sapiens] | gi\|6912486 | 15 kDa | 5 | 4 | 0 |
| U6 snRNA-associated Sm-like protein LSm2 [Homo sapiens] | gi\|10863977 | 11 kDa | 4 | 4 | 0 |
| U6 snRNA-associated Sm-like protein LSm6 [Homo sapiens] | gi\|5901998 | 9 kDa | 2 | 2 | 0 |
| U6 snRNA-associated Sm-like protein LSm3 [Homo sapiens] | gi\|7657315 | 12 kDa | 2 | 2 | 0 |
| U6 snRNA-associated Sm-like protein LSm7 [Homo sapiens] | gi\|7706423 | 12 kDa | 2 | 0 | 0 |
| U7 snRNA-associated Sm-like protein LSm11 [Homo sapiens] | gi\|27735089 | 40 kDa | 2 | 0 | 0 |
| protein arginine N-methyltransferase 5 isoform a [Homo sapiens] | gi\|20070220 | 73 kDa | 30 | 31 | 3 |
| methylosome protein 50 [Homo sapiens] | gi\|13129110 | 37 kDa | 12 | 10 | 0 |
| methylosome subunit pICln [Homo sapiens] | gi\|4502891 | 26 kDa | 7 | 8 | 0 |
| CD2 antigen cytoplasmic tail-binding protein 2 [Homo sapiens] | gi\|5174409 | 38 kDa | 13 | 12 | 0 |
| PERQ amino acid-rich with GYF domain-containing protein 2 isoform c [Homo sapiens] | gi\|156766047 | 149 kDa | 2 | 4 | 0 |
| WW domain-binding protein 4 [Homo sapiens] | gi\|6005948 | 43 kDa | 3 | 0 | 0 |
| formin binding protein 4, isoform CRA_a [Homo sapiens] | gi\|119588290 | 90 kDa | 2 | 2 | 0 |
| neurochondrin isoform 2 [Homo sapiens] | gi\|62526031 | 79 kDa | 5 | 0 | 0 |
| nuclear receptor coactivator 6 interacting protein, isoform CRA_b [Homo sapiens] | gi\|119607168 | 97 kDa | 13 | 0 | 0 |
| 7SK snRNA methylphosphate capping enzyme isoform A [Homo sapiens] | gi\|47271406 | 74 kDa | 13 | 0 | 0 |
| RNA-binding protein 40 [Homo sapiens] | gi\|40538732 | 59 kDa | 13 | 0 | 0 |

**Figure S3:**
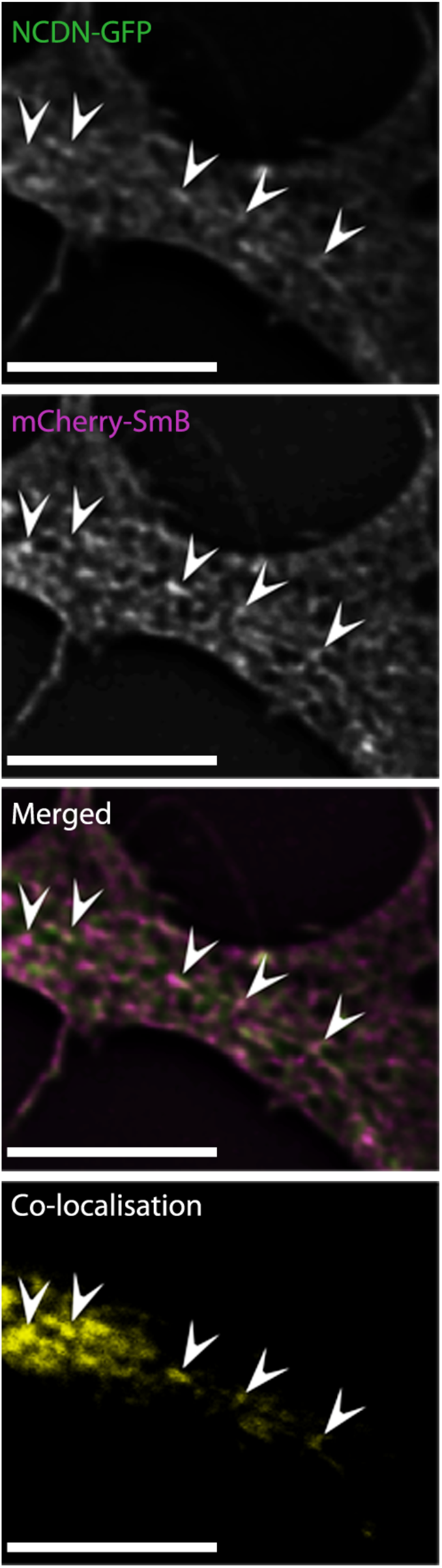
*NCDN and SmB co-localise in neurites of SH-SY5Y cells constitutively expressing mCherry-SmB, and transiently expressing NCDN-GFP. Arrows identify SmB containing-vesicles with NCDN co-localisation. Bar= 7μm, images are single deconvolved z-sections. Co-localisation images were generated by Volocity, using automatic thresholds on undeconvolved z-sections (see materials and methods).*

**Figure S4:**
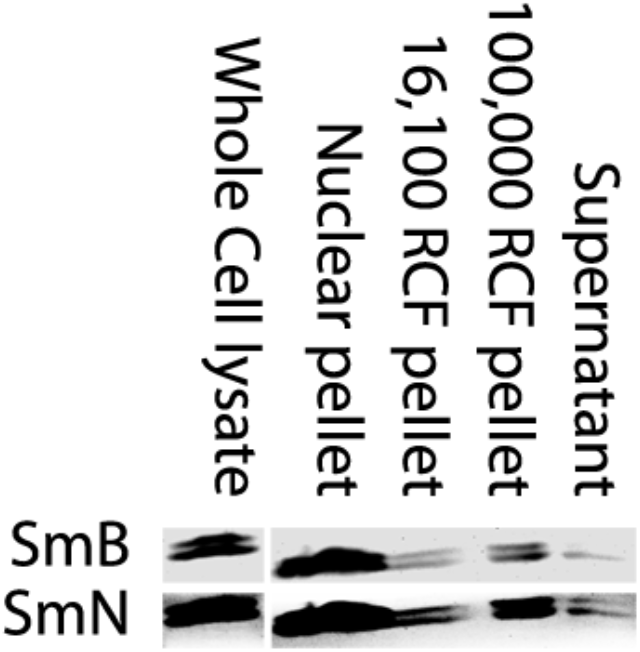
*Endogenous SmB and SmN fractionate similarly to YFP-SmB and YFP-SmN, with both present in the 100,000 RCF vesicle fraction in addition to being highly enriched in the nuclear pellet of fractionated SH-SY5Y cells. The gap between bands in the whole cell lysate and cell fractions signifies omitted lanes.*

Movie S1: punctate structures containing mCherry-SmN are mobile structures within the cytoplasm of SH-SH5Y cells. Images were taken every ~2 seconds for ~150 seconds. Movie is a projection of 3 deconvolved z-stacks taken with 0.5μm spacing. Bar=7μm.

